# Learning to Estimate Sample-specific Transcriptional Networks for 7000 Tumors

**DOI:** 10.1101/2023.12.01.569658

**Authors:** Caleb N. Ellington, Benjamin J. Lengerich, Thomas B.K. Watkins, Jiekun Yang, Abhinav Adduri, Sazan Mahbub, Hanxi Xiao, Manolis Kellis, Eric P. Xing

## Abstract

Cancers are shaped by somatic mutations, microenvironment, and patient background, each altering gene expression and regulation in complex ways, resulting in heterogeneous cellular states and dynamics. Inferring gene regulatory networks (GRNs) from expression data can help characterize this regulation-driven heterogeneity, but network inference requires many statistical samples, limiting GRNs to cluster-level analyses that ignore intra-cluster heterogeneity. We propose to move beyond coarse analyses of pre-defined subgroups by using *contextualized* learning, a multi-task learning paradigm that uses multi-view contexts including phenotypic, molecular, and environmental information to infer personalized models. With sample-specific contexts, contextualization enables sample-specific models and even generalizes at test time to predict network models for entirely unseen contexts. We unify three network model classes (Correlation, Markov, Neighborhood Selection) and estimate context-specific GRNs for 7997 tumors across 25 tumor types, using copy number and driver mutation profiles, tumor microenvironment, and patient demographics as model context. Our generative modeling approach allows us to predict GRNs for unseen tumor types based on a pan-cancer model of how somatic mutations affect gene regulation. Finally, contextualized networks enable GRN-based precision oncology by providing a structured view of expression dynamics at sample-specific resolution, explaining known biomarkers in terms of network-mediated effects and leading to novel subtypings that improve survival prognosis. We provide a SKLearn-style Python package https://contextualized.ml for learning and analyzing contextualized models, as well as interactive plotting tools for pan-cancer data exploration at https://github.com/cnellington/CancerContextualized.

**Significance Statement:** Network estimation is essential for understanding the structure and function of biological systems, but current statistical approaches fail to capture inter-subject heterogeneity or cross-modality information flow, both of which are needed for understanding complex phenotypes and pathologies. We introduce contextualized network inference, leveraging multi-view contextual metadata to capture similarities and differences among heterogeneous observations during network estimation. Sharing information across contexts enables inference at sample-specific resolution, thus quantifying variation between subjects and revealing context-specific network rewiring. Applied to tumor-specific transcriptional network inference using clinical, molecular, and multi-omic data, contextualized networks improve accuracy, generalize to unseen cancer types, and discover novel prognostic tumor subtypes. By tailoring disease models to each sample, contextualized networks promise to enable precision medicine at unprecedented resolution.

---

Tumors are heterogeneous, developing through clonal evolution that accumulates mutations, including cancer-driving single-nucleotide variants (SNVs) and somatic copy number alterations (SCNAs). In addition to tumor cell-intrinsic changes, tumors develop in and are shaped by a microenvironment that includes immune cells, the extracellular matrix, blood vessels, and surrounding cells. This extensive heterogeneity necessitates heterogeneous treatments targeted to individual patients. However, estimating treatment effects and patient prognosis at patient-specific resolution implies an n-of-1 approach to treatment that is technically and temporally infeasible. Instead, methods have historically sought to identify prognostic biomarkers that stratify patients into tumor subtype cohorts, and predictive biomarkers that identify patients who often respond to treatment. The Cancer Genome Atlas^*^ (TCGA) derives prognostic subtypes via cluster analysis on clinical and molecular data, including cancer-driving SNVs, SCNAs, DNA methylation, mitochondrial DNA, RNA-seq, miRNA, protein abundance arrays, histology images, patient demographics, and/or immunological data, and further identifies prognostic biomarkers as features that differentiate these clusters (1–25). While clusters can be analyzed in terms of feature stratification, clustering ignores the latent feature interactions and hierarchical feature relationships that define biological systems. Biomarkers identified by cluster analysis have no mechanistic interpretation and require further experimentation to validate their role in tumorigenesis and tumor pathology. Consequently, the identification of biomarkers using somatic DNA alterations or gene expression patterns has proved challenging (26). Addressing the shortcomings of cluster analysis, we focus on three questions: (1) how do we model the mechanisms of molecular interactions as they relate to tumorigenesis and treatment efficacy, (2) how do we identify prognostic biomarkers for rare diseases and outlier patients that are too sparsely sampled to cluster, and (3) how can we quantify the heterogeneity of tumor pathology, which is widely acknowledged but poorly understood, and utilize multi-view phenotypic, molecular, and environmental data to understand the forces driving heterogeneity?

GRNs help us to investigate these questions simultaneously. GRNs represent cellular circuitry, both responding to biomolecular stimulus and driving tumorigenesis. Interactions between disparate biomolecular entities can be identified at the cellular level through transcriptomic regulation, both directly and indirectly. In theory, tumor-specific GRNs capture regulatory redundancy and fragility in individual cancers. Relating tumor-specific GRNs to phenotypic, environmental, and multi-omic features can reveal how these features relate to tumor pathology and the robustness of therapeutic targets GRN restructuring and reorganization. Single-cell and multi-omic profiling have advanced the potential for studying highly context-specific regulatory relationships in GRNs, but computational methods of inferring GRNs continue to rely on partitioning samples into homogeneous sets of samples (27–30). Partition-based modeling is insufficient to capture high-resolution or continuously rewiring GRNs, which is a problem for precision oncology because some types of cancer neither form discrete clusters (31) nor cluster by tissue of origin (32).

More generally, the increase of dataset complexity, heterogeneity, and size, has motivated the development of methods of “personalized” models across several application areas (33–36). “Personalized” models seek to represent heterogeneous distributions as sample-specific distributions *X*_*i*_ ∼ *P*_*i*_ (*X*) , where *i* indexes a sample *X*_*i*_ and *P*_*i*_ corresponds to the sample-specific distribution. In the most difficult case of sample-specific inference, each *P*_*i*_ is observed only a single time and hence information must be shared across samples.

Toward this aim of sharing information across samples, most personalized models make the simplifying assumption that all *P*_*i*_ belong to the same family; i.e. *X*_*i*_ ∼ *P* ( *X* ∣ *θ*_*i*_) . Through this lens of personalized modeling, understanding sample heterogeneity is reframed as estimating data distributions with sample-specific parameters. Some methods provide sample-specific estimators without additional information by imposing strong biological priors (37) or using a sample-left-out approach (38, 39), but these lack desirable properties such as the ability to generalize to new samples or even test model performance on held-out data. Due to the difficulty of estimating sample-specific parameters, most methods make use of side information (e.g. sample metadata) as a contextual representation of sample-to-sample variation (40–44). Given observations *X* and contextual metadata *C*, we have

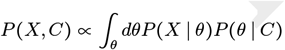

where *P* (*X* ∣ *θ*) defines the context-specific model, and *P* (*θ* ∣ *C* ) defines a context-specific density of model parameters *θ*, which we call the context encoder. One of the earliest ways to apply context encoding toward sample-specific parameter inference was the linear varying-coefficient (VC) model (44) in which linear regression parameters are predicted from context using a learned linear mapping or kernel density estimator. Extensions of this regime have been widespread (40–42, 45), but typically focus on allowing models to vary over only a few continuous covariates (44–46), or a small number of groups (40–42)

Contextualized modeling (47, 48), combines the adaptability of VC models with the power of modern deep learning architectures by implementing the context encoder as a Dirac delta distribution defined by a deterministic deep neural network *f* ,

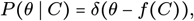

thus benefiting from a wide range of architectures targeting high-dimensional and complex data types. When contexts are unique to each sample, the inferred models are sample-specific.

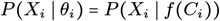

The contextualization framework also introduces the concept of model archetypes to combat the high-variance and uninterpretability of neural networks (Figure 1d). All sample-specific models are spanned by the set of model archetypes, constraining and explaining their variation through the context encoding which parameterizes this space (See Methods). These archetypes, also learned from data, link the heterogeneity of sample-specific models to variation in the context encoding and enable the sharing of information between sample-specific model inference tasks. This framework has enjoyed success is estimating heterogeneous linear effects (47, 49–51), but contextualized models have yet to be extended to the more general graphical modeling regime.

**Fig. 1.**
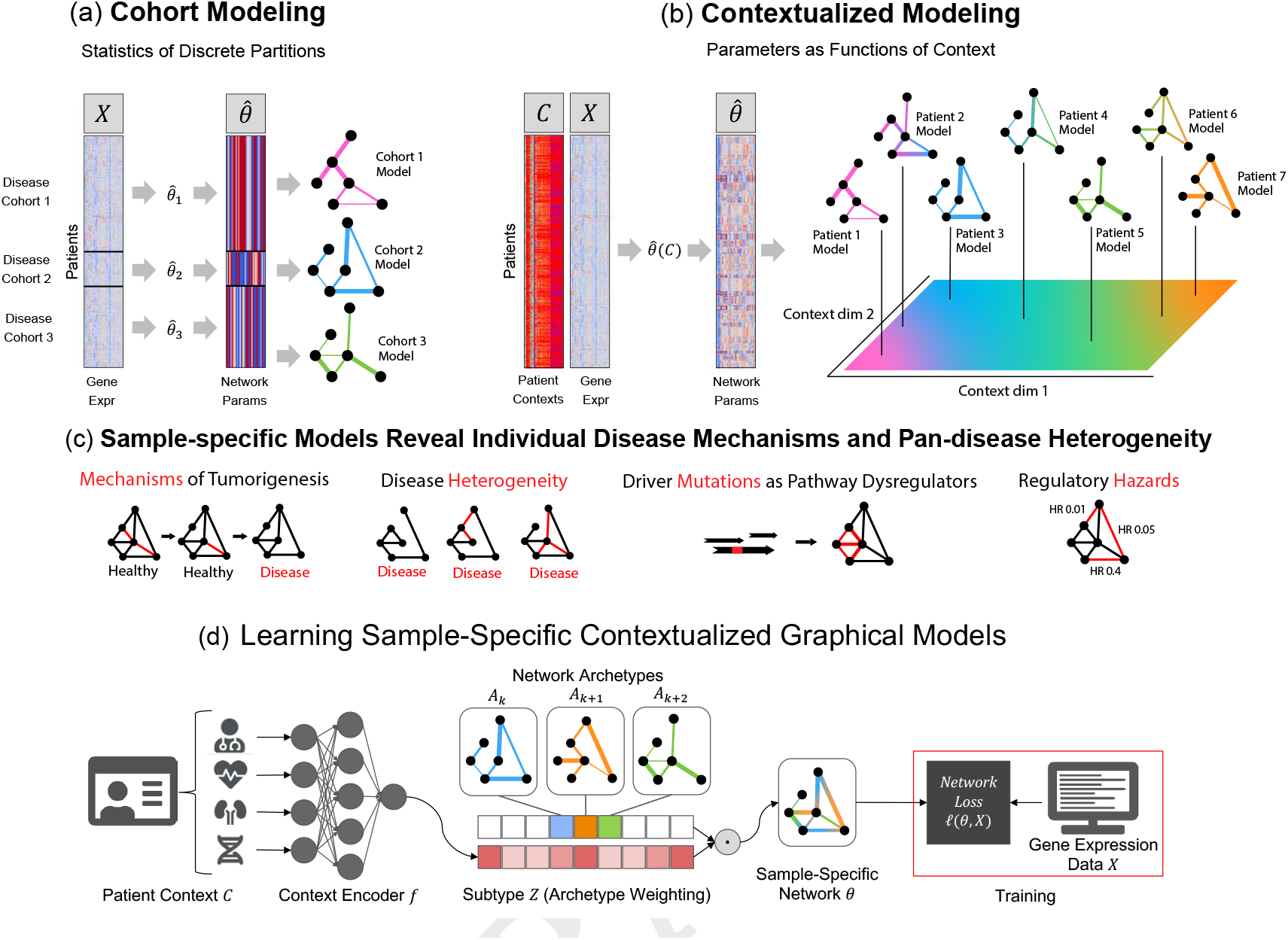
(a) Traditional modeling approaches assume each training cohort or (sub)population is homogeneous and samples are identically distributed. Cohorts must be large enough to allow robust inference, presenting a tradeoff between personalization and power. (b) Contextualization assumes model parameters are a function of context, allowing powerful context-specific inference without *a priori* clustering of subpopulations or assuming homogeneity. Contexts can be unique to each sample, permitting sample-specific model inference. (c) Sample-specific models reveal population heterogeneity, relate rare pathological mechanisms to more common ones, and provide new data views for prognosis and biomarker identification. (d) Graphical depiction of the deep learning framework. Sample context is used to predict weights on each of the model archetypes, which we call the subtype. The sample-specific network is estimated as the tensor dot product of archetypal networks and subtype weights. The network archetypes are learned simultaneously alongside the context encoder using backpropagation.

To infer tumor-specific GRNs that account for patient-to-patient heterogeneity, we propose to reframe GRN inference within the contextualized modeling paradigm (48), thereby sharing information among tumor-specific inference tasks by relating these tasks through their clinical and molecular contexts (Fig. 1). By recasting networks as the output of a learnable context encoder *P* ( *θ*∣ *C* ), our approach shares statistical power between samples while also permitting fine-grained variation to capture the complexity of sample-specific contexts such as tissue-of-origin, somatic mutation landscape, tumor microenvironment, and clinical measurements. We formulate three differentiable objectives for three types of GRNs (Markov, Neighborhood, and Correlation networks) under the contextualized modeling paradigm, and estimate sample-specific GRNs which enable sample-specific analyses of latent regulatory processes. We apply this computational framework to 7997 tissue samples from TCGA, using bulk gene expression data as network samples *X*, and immune cell infiltration metrics, patient demographics, and cancer-driver SCNAs and SNVs as context *C*. We find that contextualized networks improve prediction of held-out expression data and reveal latent heterogeneity which has previously been obscured by partition-based methods of network inference.

## Results

We introduce contextualized networks, which learn to personalize parametric network models based on context. This approach to network modeling enjoys two benefits over traditional network estimators: it scales across contexts to improve the accuracy of all networks as new contexts are included, and it allows the incorporation of multi-view context information to personalize the model. In studies on both simulated and real data, contextualized networks achieve high accuracy with as few as one training sample per context, while also generalizing to entirely unseen contexts. On real data, we apply contextualized networks to infer tumor-specific GRNs for 7997 tumors, which learn to model the effect of individual clinical and molecular contexts on GRN structure and parameters, revealing latent GRN-based drivers of GRN dysregulation and tumor heterogeneity. We evaluate our 7997 tumor-specific GRNs for new clinical and biological insights, discovering robust state-of-the-art prognostic subtypes for thyroid carcinoma. Finally, patient-specific networks relate prognostic biomarkers to changes in specific regulatory modules and gross GRN organization, and identify candidate biomarkers for further investigation.

### Unification of Markov, Correlation, and Neighborhood Network Objectives

Statistical models for GRN inference can often be categorized as variants of four probabilistic models: Markov networks, which represent pairwise dependencies, Pearson’s correlation networks, which represent pairwise correlations, neighborhood selection networks, which represent each node as a linear combination of its neighbors, and Bayesian networks, and which represent directed and acyclic interactions. We focus on Markov, correlation, and neighborhood networks, unifying these models through linear reparameterization (see Methods), thus enabling them to be contextualized uniformly with no change to the underlying contextualization framework.

Furthermore, linear parameterization gives a differentiable objective for optimizing each model, where the linear residual errors define mean-squared errors (MSEs) for measuring goodness-of-fit, which are also proportional to the negative log-likelihood of the data under the chosen network model (see Methods). Beyond the modularity of this approach and its alignment with gradient-based optimization methods, MSEs enable benchmarking model performance in the absence of gold-standard networks with known structures. Thus, we enable quantitative comparison against baseline methods for network modeling even when tumor networks are too heterogeneous and individualized to determine gold-standard network structures.

### Simulations

The convergence rate of traditional network estimators is based on the number of experimental replicates, i.e. independent and identically distributed (i.i.d.) data draws. However, scaling within an i.i.d. domain is orthogonal to the goals of sample-specific modeling in heterogeneous regimes. Contextualization addresses this by providing a new mechanism to scale model performance. In addition to the traditional route of increasing domain-specific or context-specific data collection, contextualization also scales performance with the total contexts available to the model, improving with the addition of new experimental conditions. Arranged as axes, we describe these as “vertical” and “horizontal” scaling, respectively. We compare vertical and horizontal scaling properties of contextualized, context-grouped, and population model estimators by simulating known networks. Specifically, we simulate a context-varying Gaussian, which provides a convenient testing ground our Gaussian-based network models.

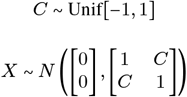

Contextualized models are the only method capable of horizontal scaling, which drastically reduces the burden of vertical scaling (Fig. 2). With sufficient horizontal scaling, contextualized networks remain accurate even in sample-specific regimes and converge to true network parameters by learning to relate heterogeneous data through contextual metadata. However, with insufficient horizontal scaling, contextualized networks can be inferior to context-grouped or population modeling methods. For data domains with sample scarcity, contextualization presents a new approach for improving model performance by sharing information across different contexts.

**Fig. 2.**
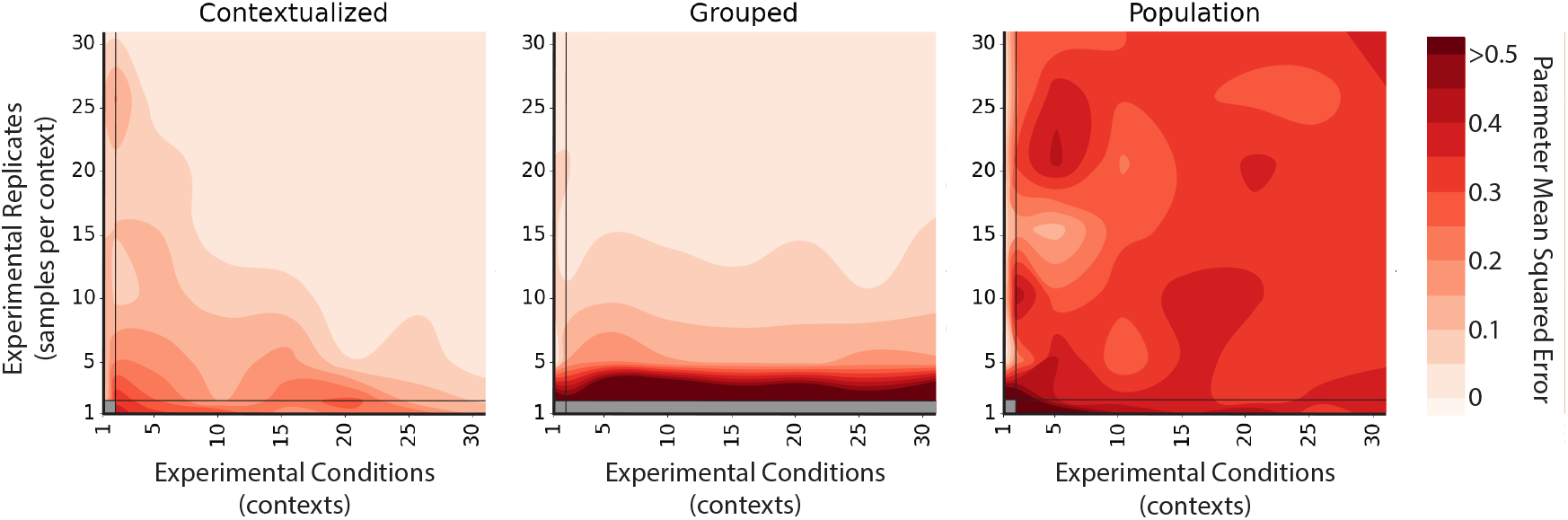
Error contours of Markov network estimators measured across experimental conditions (contexts) and experimental replicates (samples per context). Traditional estimators only scale vertically, improving error with experimental replicates or i.i.d. data draws. Modeling with heterogeneous or observational data requires estimators to scale horizontally, improving with more conditions or contexts. Contextualized modeling achieves this by learning to encode contextual information into model parameters. Population estimates a single model for all contexts, Grouped estimates a model for each context separately. Parameter mean-squared error is taken between the predicted and ground truth precision matrices of the Markov networks, and averaged over five bootstrapped runs.

### Contextualized Networks Improve Likelihood of Held-Out Expression Profiles

Contextualization improves the fit of network models to gene expression data (Table 1). We benchmark the contextualized networks by comparing them against several granularities of partition-based models: (1) a population network model that estimates the same network for all samples, (2) cluster-specific networks that are estimated independently for each cluster of contextual information, and (3) disease-specific networks that are estimated independently for each cancer type. For all three network models, we evaluate the fit of the network model to actual expression data. These predictive performances are measured as MSEs between predicted and observed expression data, a convenient result of our linearization and unification of correlation, Markov, and neighborhood selection objectives (see Methods). Relative to disease-specific model inference (the best baseline method), contextualized networks reduce modeling error on average by 14.6% for Markov networks, 18.1% for neighborhood selection, and 20.2% for correlation networks. Contextualized networks achieve this improved predictive performance by accounting for contextual dependencies in model parameters without imposing prior assumptions on the form of these dependencies. As a result, contextualized graphical models capture highly localized and context-specific effects that can be overlooked by group-level modeling approaches (e.g. cluster-specific, disease-specific models).

**Table 1.** Error of inferred networks on pan-cancer data. For all three types of networks (Markov, Neighborhood, and Correlation), we report the mean squared error (MSE) of each network versus observed gene expression (See Methods). Reported values are mean ± std over 30 runs with a bootstrapped training set and randomly initialized model weights. Error reduction is reported relative to the best baseline, which in all cases is disease-specific modeling.

|  | Markov | Neighborhood | Correlation |
| --- | --- | --- | --- |
| Population | $0.985 \pm 0.006$ | $0.984 \pm 0.004$ | $0.963 \pm 0.000$ |
| Cluster-specific | $0.365 \pm 0.014$ | $0.349 \pm 0.012$ | $0.683 \pm 0.052$ |
| Disease-specific | $0.368 \pm 0.003$ | $0.351 \pm 0.003$ | $0.673 \pm 0.002$ |
| **Contextualized** | **$0.322 \pm 0.014$** | **$0.296 \pm 0.013$** | **$0.529 \pm 0.019$** |
| Error Reduction | $14.6\% \pm 3.4\%$ | $18.1\% \pm 3.3\%$ | $20.2\% \pm 3.4\%$ |

### Contextualized Networks Share Power Between All Cancer Types and Infer Models for Unseen Diseases

Contextualization relates transcriptional regulation to genomic variation through a context encoder. During training, the encoder learns to modify the parameters of a downstream network model in response to contextual signals. At test time, the encoder uses learned context signals to generalize between sparsely sampled contexts. Splitting model performance by disease, contextualization sets a new state-of-the-art on 22 of 25 disease types (Fig. 3a). Rare or undersampled diseases like kidney chromophobe and glioblastoma multiforme can especially benefit from contextual signals learned from well-sampled diseases in similar tissues. In disease-specific modeling, these smaller subpopulations must either be lumped within a larger tissue group, ignoring subpopulation heterogeneity, or modeled individually, sacrificing statistical power in a “large *p* small *n*” regime. For example, there are *n =* 75 training samples from kidney chromophobe patients, while each disease-specific network has 50 × 50 edges, or *p =* 2500 parameters; estimating a disease-specific network from such limited data would be prohibitively high-variance for disease-specific modeling but is straightforward for contextualized networks.

**Fig. 3.**
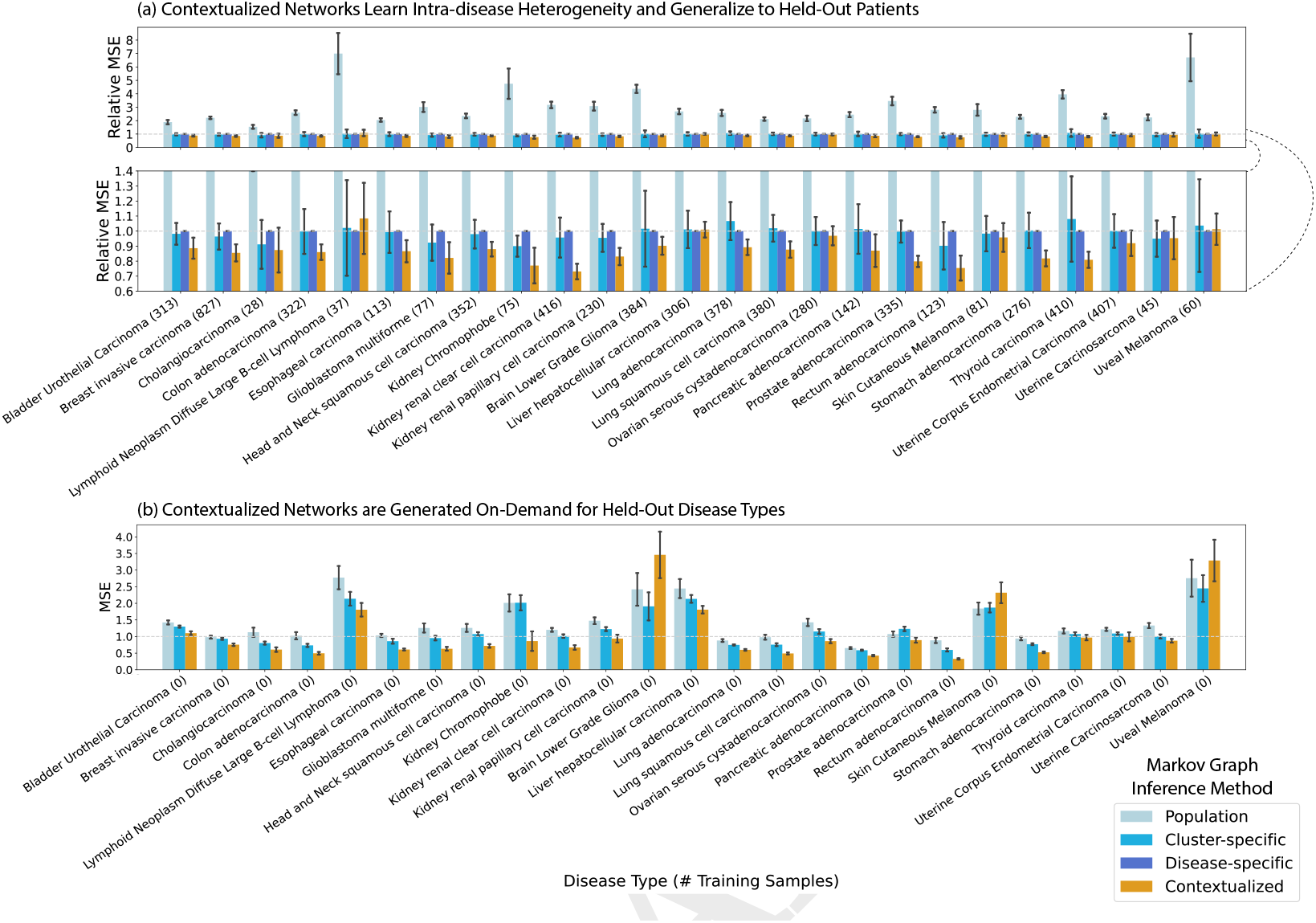
Performance of Contextualized Markov Networks broken down by disease type. **(a)** Testing on a random split of held-out patients. Mean-squared error (MSE) for Markov networks is defined in the Methods. Relative MSE scales the MSEs of all models against the Disease-specific MSEs. **(b)** Disease-fold cross-validation, in which each the 25 disease types is held out from training and evaluated only at testing time. We evaluate in terms of absolute MSEs, as Disease-specific network inference cannot be applied in this regime. Results are from 30 bootstrapped runs for each hold-out disease type and the hold-out patient set. Bar height is the disease-averaged error metric of the bootstrap-averaged network models. Error bars are the standard deviation over bootstraps of the disease-averaged error metric of the network models.

Furthermore, contextualization adapts models to unseen contexts at test time, responding to even extreme distribution shift (Fig. 3b). For completely unseen contexts, the context encoder can still leverage learned relationships between contexts and models to infer *zero-shot* network models on demand. We evaluate model performance through a disease-fold cross validation, where we hold out each of the 25 disease types in turn and learn to contextualize networks on the remaining 24. Notably, disease-specific modeling cannot be applied in this regime. In contrast, contextualized networks improve model performance and reduce error on 22 of 25 hold-out diseases, even when generalizing to an entirely new disease type.

### Contextualized Networks Reveal Tissue-Specific Regulatory Modules

Contextualization produces context-specific network models, resulting in patient-specific networks for all 7997 patients in our TCGA dataset. Organizing patients according to their network models reveals that tissue type is a primary driver, but not the sole factor in determining gene-gene interactions (Fig. 4). In particular, diseased networks differ drastically from healthy networks, while gene and PCA-derived metagene expression profiles are still largely tissue-derived. Additionally, intra-disease and inter-disease disease subtypes are visible even at pan-cancer resolution, making obvious common tumorigenesis mechanisms that underly noisy gene expression dynamics. These subtypes are further explored in Figure 5 and in Supporting Information. We also provide tools for on-demand and interactive plotting from population-level to disease-level to sample-specific at https://github.com/cnellington/CancerContextualized.

**Fig. 4.**
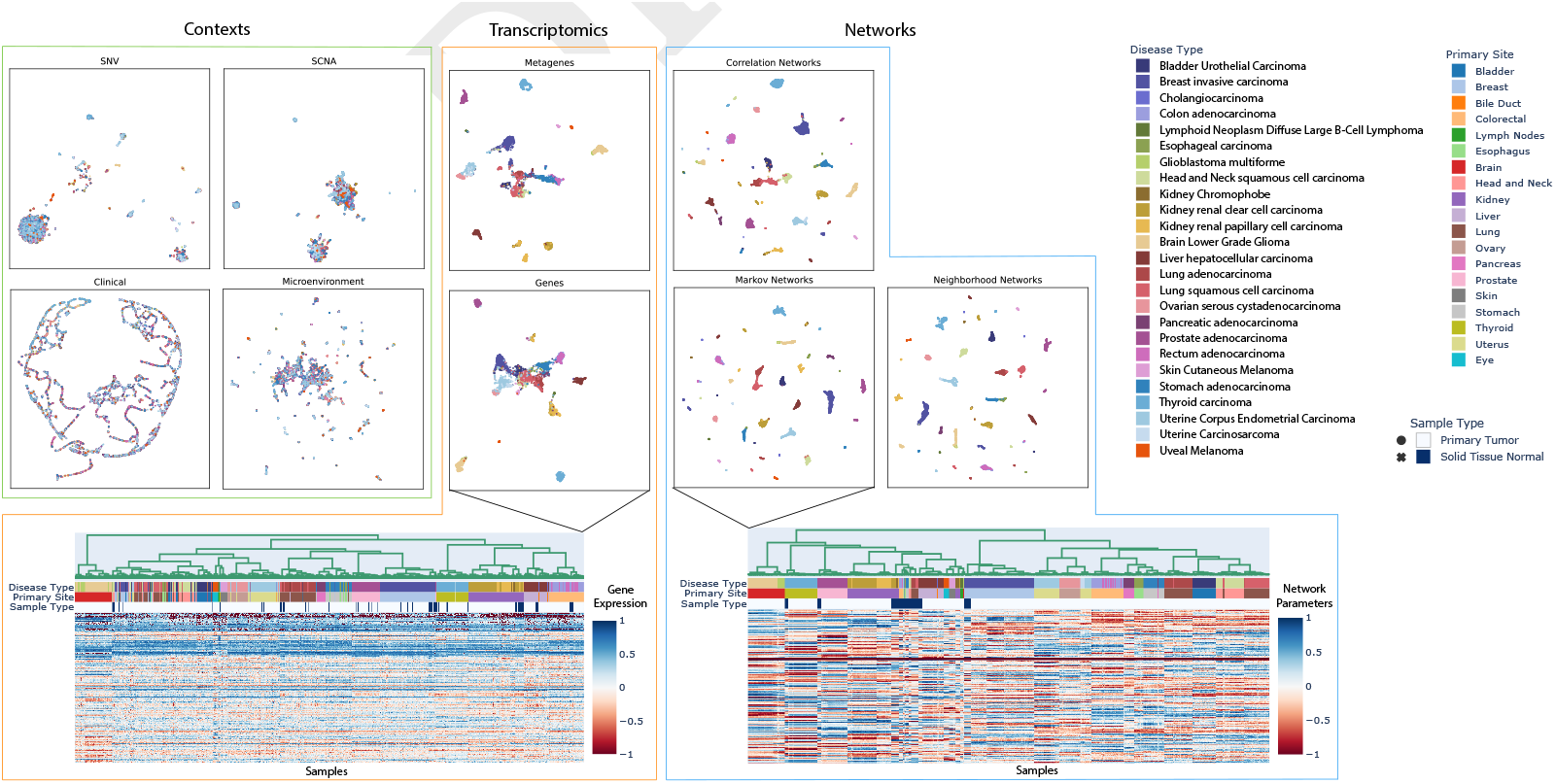
UMAP embeddings, colored by disease type, reveal the organization of different data views with respect to known disease types. Context views are used as input for the context encoder. Transcriptomic views recapitulate disease types, relating to known cell-of-origin patterns (32). Contextualized networks reorganize patients to refine and separate disease types into subtypes based on tumor-specific GRNs. Refined network-based subtypes are further explored in Figure 5 and the Supporting Information.

**Fig. 5.**
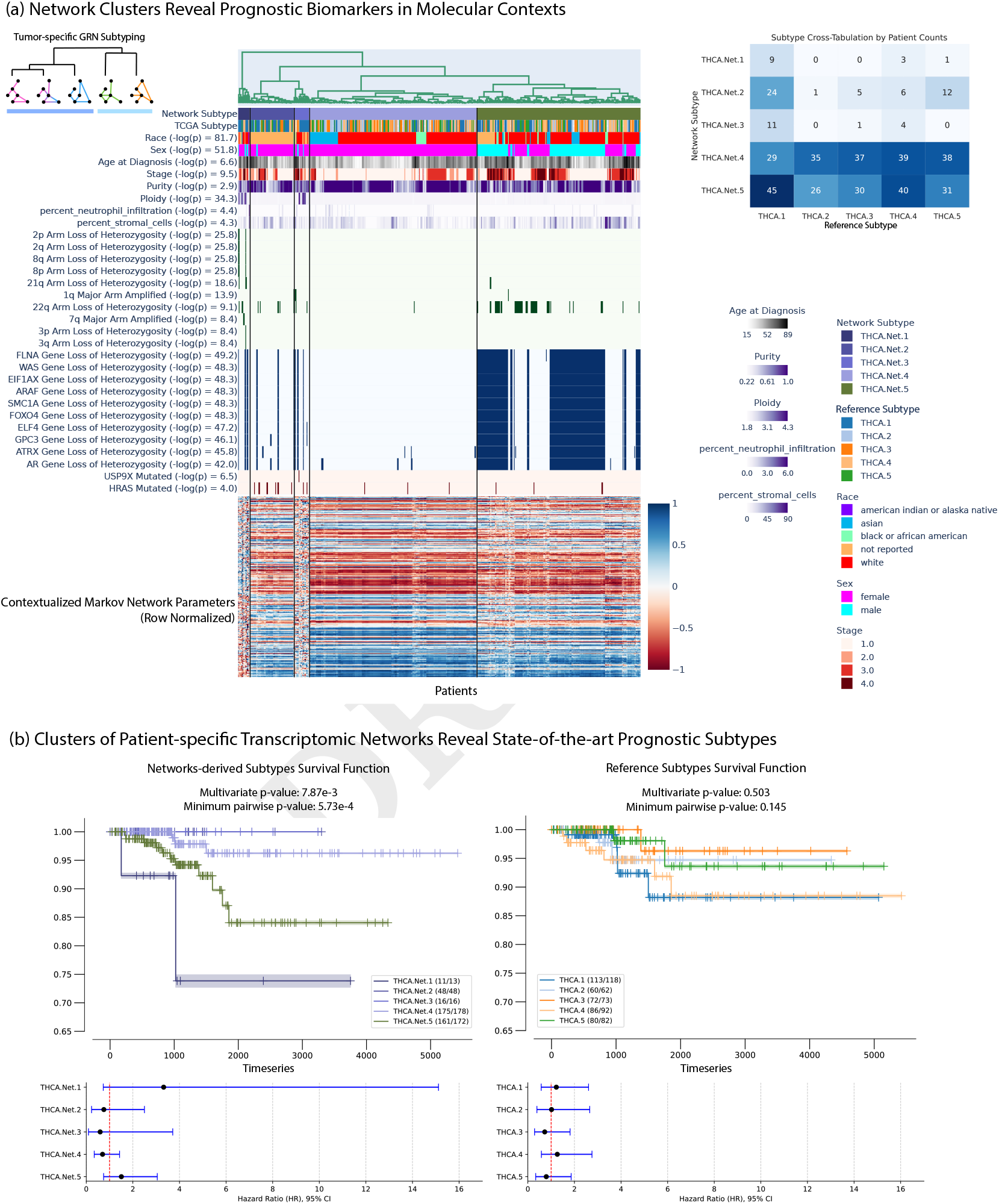
Exploration of network subtypes for thyroid carcinoma (a) looking at correlated clinical information, arm-level copy alterations, gene-level copy alterations, and gene-level single nucleotide variations, and (b) comparing against state-of-the-art reference subtypes (52). An interactive version of this plot can be produced on-demand with tools at https://github.com/cnellington/CancerContextualized, and re-plotted for other diseases or cohorts.

### Contextualized Networks Discover Novel Prognostic Subtypes for Thyroid Carcinoma

Contextualized GRNs discover the first significantly prognostic molecular subtypes for thyroid carcinomas (THCAs) (Fig. 5). We produce GRN-based subtypes by clustering samples according to the sample-specific parameters of Contextualized GRNs. To benchmark their prognostic ability, we compare our GRN-based subtypes against state-of-the-art reference subtypes (52), evaluating the survival splits of both subtyping methods while ensuring the number of clusters are the same (Fig. 5b). Previous state-of-the-art molecular subtypes are highly consistent across studies, but not prognostic (52, 53). Contextualized GRN-based subtyping reveals several clinically meaningful subtypes: one with extremely poor survival prognosis (THCA.Net.1) and two with no recorded deaths (THCA.Net.2 and THCA.Net.3). These three GRN-based subtypes stratify the previous state-of-the-art “RAS-like” molecular subtype (52). Contextualization allows us to relate these GRN clusters to clinical and molecular contexts associated with each subtype (Fig. 5a). THCA.Net.1’s poor prognosis is defined by chromosomal instability and tumor ploidy. THCA.Net.2 and THCA.Net.3 both show enrichment for HRAS mutations. THCA.Net.2 and THCA.Net.3 are mainly differentiated differentiated by patient demographic reports, with THCA.Net.2 containing almost exclusively patients with no race reported. This split, as well as the race and sex-related subgroups in THCA.Net.4 and THCA.Net.5, are supported by known gender and race disparities related to thyroid cancer presentation (54, 55). Contextualized networks combine both clinical and molecular data sources toward a cohesive molecular representation of tumor state, and relate the resulting tumor-specific GRNs back to contexts to identify stratifying biomarkers in all contextual data types.

### Contextualized Networks Unify Pan-cancer Subtypes and Improve Survival Prediction

We repeat the same procedure for all 25 tumor types in our dataset and compare against reference molecular subtypes, keeping the number of clusters matched to each study (1–24). For diseases where no reference subtypes exist or can be mapped to our dataset, we select the number of networks subtypes based on silhouette score of the network subtype clusters. We find that network-based subtypes are more prognostic on average than both expression-derived subtypes and state-of-the-art reference subtypes (Table 2). Previous molecular subtypes also exclude demographic and immune data, which contextualization naturally incorporates alongside molecular features, learning to relate these disparate feature sets as they relate to GRN restructuring. Subtype comparisons by disease can be plotted on-demand with tools at https://github.com/cnellington/CancerContextualized. In the majority of other tumor types, contextualized modeling does not identify sex or race as significant factors driving GRN variation. However, the ones that do include breast invasive carcinoma, esophageal carcinoma, and kidney renal clear cell carcinoma, which have known race and sex disparities (25, 56, 57).

**Table 2.** Stratification disease subtypes in terms of survival. Survival tests quantify the difference in survival distributions between groups as a p-value. Contextualized networks improve on both tests on average by several orders of magnitude compared with other subtyping methods. The multivariate log-rank test quantifies overall stratification of survival distributions across all subtypes. The minimum pairwise result is the minimum p-value of all pairwise subtype tests, showing the maximum survival stratification between prognostic subtypes.

| Average -log(p) | Expression | Reference | Networks |
| --- | --- | --- | --- |
| Multivariate log-rank test | 8.53 | 9.65 | **11.24** |
| Minimum Pairwise log-rank test | 8.27 | 9.55 | **11.71** |

In addition to comparing the benefit of organizing patients using transcriptional network similarities through subtyping, we also run a survival regression based on different patient representations (Table 3). We find that tumor-specific networks lead to more accurate survival predictors than previous molecular subtypes on both a per-patient and per-tissue basis.

**Table 3.** Survival regressors trained on different patient representations evaluated in terms of concordance index on a held-out set. Expression use gene expression as input to the regressor. Reference uses curated molecular subtypes for each patient. Networks use sample-specific transcriptional networks as input to the regressor. One regressor was trained for each tissue type.

| Concordance Index | Expression | Reference | Networks |
| --- | --- | --- | --- |
| Patient Average | 0.620 | 0.598 | **0.647** |
| Tissue Average | 0.565 | 0.569 | **0.592** |

## Discussion

In spite of the evidence for functional convergence (58, 59), it is challenging with current statistical methods to identify biomarkers that define similar phenotypes in genetically diverse contexts in order to guide treatment. In this study, we propose contextualized GRNs as cohesive sample-specific representations of latent tumor states underlying disease progression and patient survival. Our models reveal new insights about cancer heterogeneity by relating transcriptomic, genetic, immune, and clinical factors to tumor regulatory network topology. In Figures 4 and 5, contextualized GRNs provide an intuitive way of identifying both subpopulations with differential transcriptomic regulation and the pathway-level modules of genes that should be studied as potential biomarkers, as well as the likely effect size of pathway dysregulation. Contextualized GRNs further identify contextual signals differentiating these subpopulations, exploiting these signals for predictive accuracy (Fig. 3) and providing new leads for traditional classes of genomic biomarkers (Fig. 5).

More broadly, contextualized modeling seeks to estimate context-specific models beyond context-specific sampling constraints that currently prohibit individualized and independent analysis of patients. By sharing information among samples while also allowing sample-specific variation, our framework models complex and dynamic distributions despite physical and technical barriers that prohibit sample-specific inference. For instance, observational patient data is often heterogeneous, suffering from complex confounders relating to environment, genetics, and individual histories. However, controlling for all conditions and contexts simultaneously leads to subpopulations with as few as just one sample – too small to infer accurate context-specific models. We explore this tradeoff in a simulation study, showing how both group-specific models and population-level models fail in heterogeneous and sample-specific regimes which require horizontal scaling across data domains (Fig. 2). Contextualized models naturally account for non-identically distributed data and even improve performance by incorporating multiple data views as contexts for model estimation, providing a principled method for performing statistical inference on heterogeneous, observational, and multi-view data.

Finally, contextualized modeling raises questions about how to interpret and apply populations of sample-specific models, which we leave partially open to future work. In this study, we show that a measure defined by model parameters can be used to traverse the sample-specific model space. Another route for future work is to interpret the archetypes themselves. In this study, archetypes serve to regularize sample-specific model generation, but this same mechanism also defines a polytope for all possible sample-specific models (60). New statistical tests are also needed to quantify the degree of heterogeneity in data and the effects of contextual features on model parameter variation (61).

## Materials and Methods

Contextualization is based on two simple concepts: a context encoder which translates sample context into model parameters, and a sample-specific model which represents the latent context-specific mechanisms of data generation. This view conveniently unifies both varying-coefficient models (44), and subpopulation and partition-based approaches, such as cluster analysis and cohort analysis (62). By learning how models change in response to context, contextualization enables powerful control over high-dimensional and continuously varying contexts, discovering dynamic latent structures underlying data generation in heterogeneous populations and permitting GRN model inference at even sample-specific resolution.

### Contextualized Networks

We seek a context-specific density of network parameters ℙ (*θ* ∣ *C* ) such that

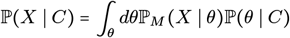

is maximized, where ℙ_*M*_ (*X* ∣ *θ*) is the probability of gene expression *X* ∈ ℝ^*p*^ under network model class *M* with parameters *θ* ∈ ℝ^*p*×*p*^, and *C* is sample context which can contain both multivariate and real features. To overcome *θ* being a high-dimensional, structured latent variable, we assume that all contextualized networks lie on a subspace spanned by a set of *K* network archetypes 𝒜 ∶= span ({*A*_*k*_ ∈ ℝ ^*p* × *p*^ ∶ *A*_1_, …, *A*_*K*_}), i.e. *θ* ∈ 𝒜. We further introduce a latent variable (“subtype”) *Z* ∈ ℝ^*K*^ which coordinates the archetypes such that

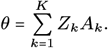

The context-specific network model *θ*, and subsequently the gene expression observations *X*, are also assumed independent of context given *Z*, i.e. *C* ⊥ ( *X, θ*) ∣ *Z*. Finally, to enable efficient gradient-based optimization, we assume *Z* is a deterministic function of context *Z* = *f* (*C*) . In this way, we constrain *θ* as a convex combination of network archetypes via latent mixing.

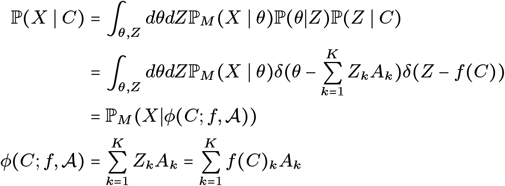

Where the context encoder *ϕ* (*C*; *f*, 𝒜) is parameterized by a differentiable context-to-subtype mapping *f* and the set of archetypes 𝒜. This architecture is shown in Figure 1d, and is learned end-to-end with backpropagation. While the archetypal networks *A*_*k*_ provide an obvious way to incorporate prior knowledge of network structures for intitialization or regularization, no prior knowledge is required. In all experiments reported here, we do not use any prior knowledge of network structure or parameters.

This framework unites three different perspectives of GRNs: (1) Correlation networks, in which network edges are the pairwise Pearson’s correlation between nodes, (2) Markov networks, in which edges are the pairwise precision values representing conditional dependencies between nodes, and (3) Neighborhood selection networks, in which edges represent directed linear relationships between nodes. The key challenge for each network class is to define a differentiable loss function 𝓁_*M*_ that is proportional to the negative log probability of our contextualized network model.

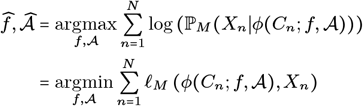

The loss objective can be used in the end-to-end optimization, solving for the context encoder and the network archetypes simultaneously, to infer the context-specific parameters *θ*. Below, we outline a unifying linear parameterization of each network loss. Implementation details are discussed in Supporting Information.

### Contextualized Neighborhood Selection

We first apply contextualization to the neighborhood selection algorithm proposed by Meinhausen and Buhlmann (63). The direct relationship of this model to lasso regression (64) links contextualized neighborhood selection to original works on contextualized linear models (47) as well as earlier works on time-varying networks (41) and tree-varying networks (42), making it a convenient stepping stone toward the graphical models in the sequel. The model is a Gaussian graphical model where *X* ∼ *N* (0, Ω −^1^) and precision matrix Ω is the inverse of the covariance matrix and has sparse off-diagonal entries. Based on an equivalence between precision, partial correlations, and multivariate regression coefficients (65, 66), we have that

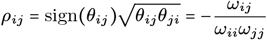

where *ρ*_*ij*_ is the correlation between features *X*_*i*_ and *X*_*j*_ conditioned on all other features *X* − {_*ij*_} , *θ*_*ij*_ is the coefficient for *X*_*j*_ in a multivariate regression onto *X*_*i*_ from all other genes *X* −*i*, and *ω*_*ij*_ are elements of the precision matrix. The dependency structure defined by the Markov random field of the model above emerges as

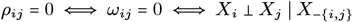

In the population setting, this dependency structure is discovered by solving the lasso regression for every feature *X*_*i*_ given every other feature *X*_−*i*_. This regression maximizes *P* (*X*_*i*_ ∣*X*_−*i*_) via the loss

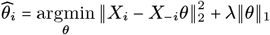

resulting in edges between *X*_*j*_ and *X*_*j*_ for every *j* ≠ *i* where *θ*_*ij*_ ≠ 0, or no edge if *θ*_*ij*_ = 0. Equivalently, we parameterize the neighborhood selection objective using the square matrix of regression parameters *θ* ∈ ℝ ^*p*×*p*^.

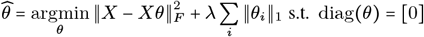

To contextualize this network objective, we replace *θ* for each sample with a context-specific *θ*_*n*_ = *ϕ*(*C n* ; *f*, A) . Finally, we define a function *ϕ*′ to mask the diagonal of *θ*, presenting the loss function 𝓁_*NN*_ for contextualized neighborhood selection networks

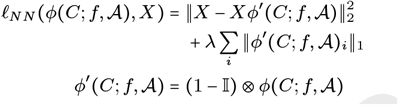

where ⊗ is the Hadamard product. In previous works on time-varying and tree-varying networks (41, 42), minimizing this loss has depended on the convexity of the objective with respect to *θ*, and subsequently the parameters *f* and 𝒜 here. While we note that this is guaranteed convex for linear *f* , in practice we utilize a neural network as our choice of *f* which are highly performant despite their non-convexity.

### Contextualized Markov Networks

Following (45), we can make further assumptions to improve the alignment of the neighborhood selection objective with the underlying Gaussian graphical model, and even recover exact precision. Assuming a constant diagonal precision *ω*_*ii*_ = *ω*_*jj*_ ∀ *i*, *j*, the neighborhood selection objective results in a proportionality between the regression and the precision matrix.

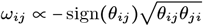

Assuming unit diagonal precisions *ω*_*ii*_ = 1, the proportionality becomes exact equivalence. Furthermore, this proportionality induces symmetry in the regression, i.e. *θ*_*ij*_ = *θ*_*ji*_. We encode this in the objective by requiring our estimate for *θ* to be a symmetrically augmented matrix based on *γ*, i.e. *θ = γ*+ *γ*^*T*^

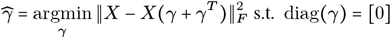

If Ω is sparse, we can again apply lasso regularization to the multivariate regression objective (63). Given the similarity between this differential Markov network objective and the neighborhood selection objective, we follow the exact contextualization procedure from above to contextualize *γ* and arrive at a loss function 𝓁_*MN*_

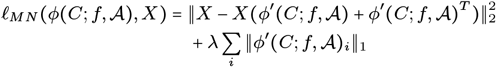

where *ϕ*^′^ is defined identically for masking the diagonal. The resulting contextualized precision matrix estimate is 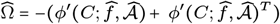. In practice, we do not threshold the estimated precision as we did in neighborhood selection. We represent the Markov network using the full precision matrix, retaining information about the dependency structure as well as the dependency strength based on the equivalence to partial correlation above.

### Contextualized Correlation Networks

Correlation networks are simple to estimate and often state-of-the-art for gene regulatory network inference (29); contextualized correlation expands this utility to the granularity of sample-specific network inferences. To estimate sample-specific correlation networks, we assume the data was drawn from *X* ∼ *N* (0, Σ) and use the well-known univariable regression view of Pearson’s marginal correlation coefficient:

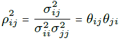

where the covariance matrix Σ has elements *σ*_*ij*_ , and 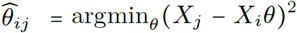. This form converts correlation into two separable univariate least-squares regressions that maximize the marginal conditional probabilities *P* (*X*_*i*_ ∣*X*_*j*_) and *P* (*X*_*j*_ ∣*X*_*i*_) . Contextualizing this differentiable objective, we get the contextualized correlation network loss

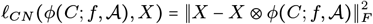

where the context-specific c orrelation matrix is reconstructed as 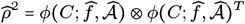.

### Baselines

We compare contextualized modeling with several traditional approaches for context-controlled and context-agnostic inference, including population modeling, cluster modeling, and cohort modeling (Fig. 6). A population model assumes that the entire sample population is identically distributed. As a result, population modeling infers a single model representing all observations. In reality, sample populations often contain two or more uniquely distributed subpopulations. If we expect that there are several subpopulations with many observations each, and that these subpopulations can be stratified by context, it may be appropriate to cluster the data by context to identify these subpopulations and then infer a model for each context-clustered subpopulation. This assumes that all context features are equally important and therefore does not tolerate noise features well. Alternatively, when subpopulation groupings are known to be determined by a few important features, cohort modeling is more appropriate. Sample cohorts can be identified based on prior knowledge about important context features (e.g. disease type).

**Fig. 6.**
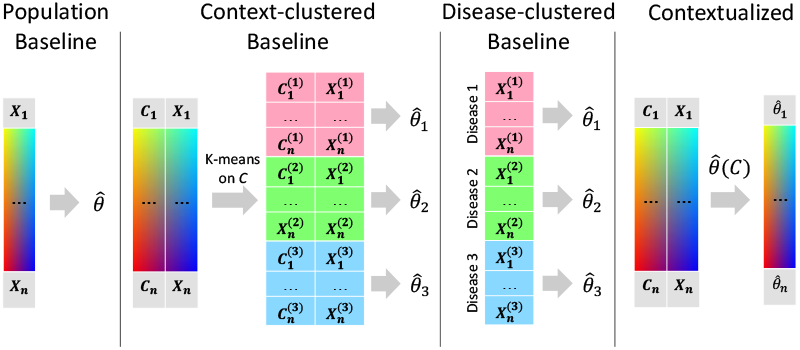
Modeling regimes for personalized inference.

The baseline modeling regimes enjoy the benefits of traditional inference methods (i.e. identifiability, convergence) by relying on the assumption there are a discrete number of subpopulations underlying the observed data that are each defined by a latent model, and each of these subpopulations is well-sampled. This assumption is rarely, if ever, satisfied in a real-world setting. We develop contextualized modeling as a synthesis between traditional statistical inference and modern deep learning to enable model-based analysis of heterogeneous real data. Contextualized modeling assumes a functional dependency between models, but unlike prior methods makes no assumption about the form or complexity of this dependency. As such, contextualized models permit context-informed inference even when contexts are sparsely sampled and high dimensional.

### Data

Our dataset is constructed from The Cancer Genome Atlas^†^ (TCGA) and related studies, covering 7997 samples from 7648 patients with 6397 samples for training and validation and 1600 as testing. For context, we use clinical information, biopsy composition, SCNAs and cancer-driving SNVs (see Supporting Information).

#### Gene Expression

We gathered samples with open access TPM-normalized expression data from TCGA. From the gene expression panel, we selected known oncogenes and tumor suppressor genes annotated by COSMIC (67). Afterward, the data was log ( *x* + 10^−3^)transformed. Finally, the data was compressed into metagenes using a PCA transformation learned on the training set. Networks were learned to model the metagene expression data.

Network models also provide a new opportunity for dealing with batch effects in expression data. Most batch-effect correction methods make strict assumptions about homogeneity between groups, which is mutually exclusive with our study design. Luckily, network optimization objectives play nicely with batch effects. If batch effects can be isolated, these can be treated as noise features where distribution shifts cannot be predicted or explained by covariate shifts or other features. Network modeling objectives learn to ignore noise features which are not predictive of others in the network. Thus, network modeling only requires batch effect isolation, not correction. PCA is convenient for isolating these effects due to the feature orthogonality and its preference for global effects over local effects. The main downside of leaving batch PCs in the metagene data is that noise features inflate all model errors by a constant amount, but this is unimportant for relative performance comparisons.

We used 50 metagenes due to hardware limitations (see Supporting Information). These 50 metagenes captured 79.47% of the variance in the pre-PCA data.

### Subtyping

To benchmark their prognostic ability, we compare our GRN-based subtypes against state-of-the-art reference subtypes gathered using TCGAbiolinks (68). Network subtypes are inferred by clustering on network parameters, where networks are organized by hierarchical clustering with ward linkage. When reference subtypes are available, the number of clusters is matched to the number of known reference subtypes for fair comparison. When reference subtypes are unavailable, the number of clusters *k* is selected by the best silhouette score from *k* = [2, 10]. We identify some contextual features as drivers of heterogeneity, having a significant association with one network subtype compared to the rest by using a two-sided t-test on the subtype vs. remaining samples. Features with universal importance within each disease type will therefore not be associated. For each feature, we take the minimum p-value from all subtype t-tests and display the most significant features. We provide an interactive demo for subtyping with network and expression data, and comparison with reference subtypes at https://github.com/cnellington/CancerContextualized.

## Code availability

All methods are available in Contextualized, an open-source SKLearn-style Python library for contextualized modeling (62). Contextualized graphical models, as well as contextualized regressors can be estimated using an intuitive import-fit-predict workflow.

~~~
from contextualized.easy import (
  ContextualizedCorrelationNetworks
)
model = ContextualizedCorrelationNetworks()
model.fit(C_train, X_train)
err = model.measure_mses(C_test, X_test)
r = model.predict_correlation(C_test)
~~~

We provide demos and tutorials for network inference at contextualized.ml. Our code for generating the figures in this manuscript is available at https://github.com/cnellington/CancerContextualized.

## Supporting Information

### Other supporting materials for this manuscript include the following

Datasets S1 to S2

## Supporting Information Text

### 1. Resources and Reproducibility

- Contextualized documentation and installation instructions https://contextualized.ml/
- Code for model training, plotting, and evaluation https://github.com/cnellington/CancerContextualized
- Preprocessed data and contextualized networks for all 7997 patients with split labels https://zenodo.org/records/14885352
- TCGA data download portal https://portal.gdc.cancer.gov/
- TCGA reference subtypes https://bioconductor.org/packages/3.21/bioc/vignettes/TCGAbiolinks/inst/doc/subtypes.html

### 2. Implementation

Our entire framework is implemented in PyTorch using the PyTorch Lightning framework within our open-source software Contextualized (1). The context encoder, network archetypes, and contextualized network models are learned simultaneously using end-to-end backpropagation of the network loss (defined in Methods).

#### Training

The context data views (B) were concatenated sample-wise to create a single context feature vector encompassing all views for each patient. Categorical features were one-hot encoded. Healthy samples were given ideal genomic profiles with no SCNAs or SNVs. Other missing values were imputed with the feature mean. The full context vectors were compressed to 200 features using a PCA learned on the training-validation split. We split our dataset into 80% training-validation and 20% testing. We created 30 bootstraps of the training-validation set and finally split into 80% training and 20% validation, resulting in a 64-16-20 split for train-validation-test.

We bootstrapped our models, sampling the train-validation split with replacement to create 30 bootstraps. For each bootstrap, we also initialized the models randomly. We trained the models using a batch-size of 10 and a learning rate of 1e-3. We used early-stopping with a patience of 5 to end training when the minimum validation loss had not improved for 5 epochs. We retain only the model with the minimum validation loss for each bootstrap.

Following this, we use the trained models to infer 30 networks (one from each bootstrap) for each sample in the original non-bootstrapped training-validation set as well as the test set. To obtain the final predicted networks for each sample, we average the network parameters over the bootstraps to get a single bootstrap-averaged network for each sample. The mean-squared errors of the network relative to the sample expression is defined in the Methods (the mean of the linear residuals in the network objectives). We report the mean error of these networks averaged over samples in the test set. Rather than bootstrapping this entire procedure again to report the standard deviation of this result, we upper-bound the uncertainty by reporting the standard deviation of the errors from the individual bootstrap models. This procedure is repeated for all of the baseline methods. All plots can be reproduced using the error files uploaded to Zenodo with the pre-trained networks.

Additionally, we tested both gradient-based optimization and off-the-shelf SKLearn solvers for the baselines where possible, applying early stopping with the gradient-based solvers to control overfitting across all methods. However, we found that the SKLearn solvers were superior when applicable, and reported the best baseline results.

#### Context Encoder

The context encoder is implemented as a multi-layer perceptron with 3 hidden layers, each 100 neurons wide with ReLU activations. Model weights are initialized as `Uniform[-0.01, 0.01]`.

The context encoder is a highly flexible component of our framework and a driving force for future work. It can be used to enforce assumptions about the relationships between contexts and models, between context features, and about the archetype space. For instance, by using a multi-layer perceptron in this study, we naturally handle colinearities in a high-dimensional feature space. Using a neural additive model (2, 3) instead of a multi-layer perceptron would provide context-feature-specific archetype weights for interpretability. Similarly, the context encoder can be implemented as a convolutional neural network for images (4) or a recurrent neural network for timezeries (5). At the context encoder head, we currently use an unconstrained output, but applying a softmax activation would require all of the sample-specific models to lie within a polytope defined by the archetypal networks.

#### Hardware Limitations

50 metagenes were chosen due to hardware limitations, as the largest number of metagenes we tried which would fit in memory for training and inference on a MacBook Air 2020 with 16G RAM in 32-bit precision. This occupied 11.4G of memory while writing 30 bootstraps * 7997 samples * 50^2^ sample-specific network parameters to disk.

### 3. Data

#### A. Data sources

The Cancer Genome Atlas^∗^ (TCGA) is a publicly-available pan-cancer datasource containing genomic, transcriptomic, and clinical profiling of tumors from dozens of landmark studies. We queried TCGA for open access samples with bulk RNA-sequencing and merged this dataset with two follow-up studies on an overlapping set of patients.

##### Somatic copy number alterations (SCNAs)

SCNAs affect a larger fraction of the genome than do any other type of somatic genetic alteration (6) and are a major driver of expression variation in cancer (7). We used copy number profiles derived from TCGA samples using ASCAT (8) from a pan-cancer study of the role of allele-specific SCNAs in cancer (9).

##### Driver single-nucleotide mutations (SNVs)

SNVs can be classified into “driver” mutations thought to provide selective growth advantage and “passenger” mutations thought to have little role in promoting cancer development. We incorporated driver SNVs from the TCGA-derived CHASMplus dataset (10)

#### B. ontext Data Views

##### Clinical information

This data view incorporates sample tissue-of-origin, race, age at diagnosis, gender, year of birth, and days to collection provided by TCGA.

##### Biopsy Composition

This data view contains the sample’s percent tumor cells, percent normal cells, percent tumor nuclei, percent monocyte infiltration, percent lymphocyte infiltration, and percent neutrophil infiltration provided by TCGA. We also incorporate expression-derived estimates of the fraction of a sample consisting of tumor cells from (9).

##### Copy Number Alterations

From ASCAT (8), we gather whole genome doubling events as well as gain and loss events for bp-specific regions of hg19 based on data from (11). We transform these gain and loss events into both arm-level and gene-level events, where arm-level events affect 85% of an entire arm in the same event, while genes-level events affect a single gene. We transform these into number of major and number of minor chromosome arms, and the number of major and minor alleles for the set of 295 genes that overlap between COSMIC (12) and MSigDB (13). For both gene and arm-level events, we create a separate indicator for loss of heterozygosity on each gene.

##### Driver Mutations

From CHASMplus (10) we gather the mutations on all COSMIC (12) oncogenes/tumor suppresor genes and binarize the presence or absence of a mutation in each gene.

#### C. aselines

We are not aware of any other scalable meta-learning, deep learning, or varying-coefficient methods to produce context-informed correlation, Markov, and neighborhood selection networks under a universal framework. State-of-the-art gene regulatory network estimators are limited to population, cohort, and cluster-based approaches tailored to a single network model class (14–16). As such, our baselines apply the network estimators in 2 under several well-known and general paradigms for improving model personalization, broadly relating to subpopulation or cluster analysis. Our population baseline provides no personalization, learning a single model for the entire population of training samples. Our context-clustered baseline takes an unsupervised approach to personalization by first doing a k-means clustering with k=25 on the aggregated context views (3) and then inferring cluster-specific networks. Our disease-clustered baseline uses a personalization oracle, grouping samples by tumor type and then inferring disease-specific networks.

### 4. Extra Results

#### A. Simulations

We perform simulations to study the scaling properties of contextualized networks over network size (Fig. S1a, S2a) and context dimensionality (Fig. S1b, S2b).

**Fig. S1.**
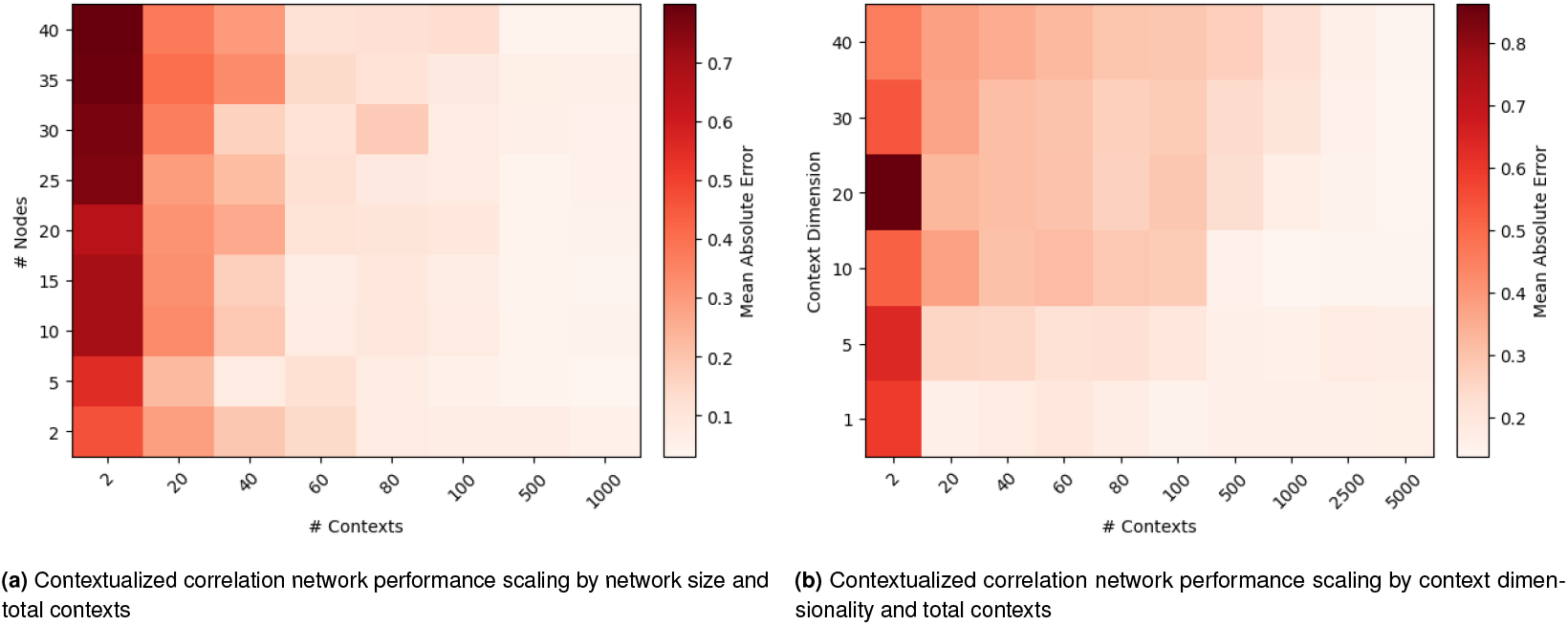
Performance of estimators on held-out data, measured across (a) network size and (b) context dimensionality (i.e. complexity of context dependence) measured as the Mean-absolute Error (MAE) between predicted and ground truth correlation networks, averaged over three bootstrapped runs. (a) Contextualized Markov network performance scaling by network size and total contexts (b) Contextualized Markov network performance scaling by context dimensionality and total contexts

**Fig. S2.**
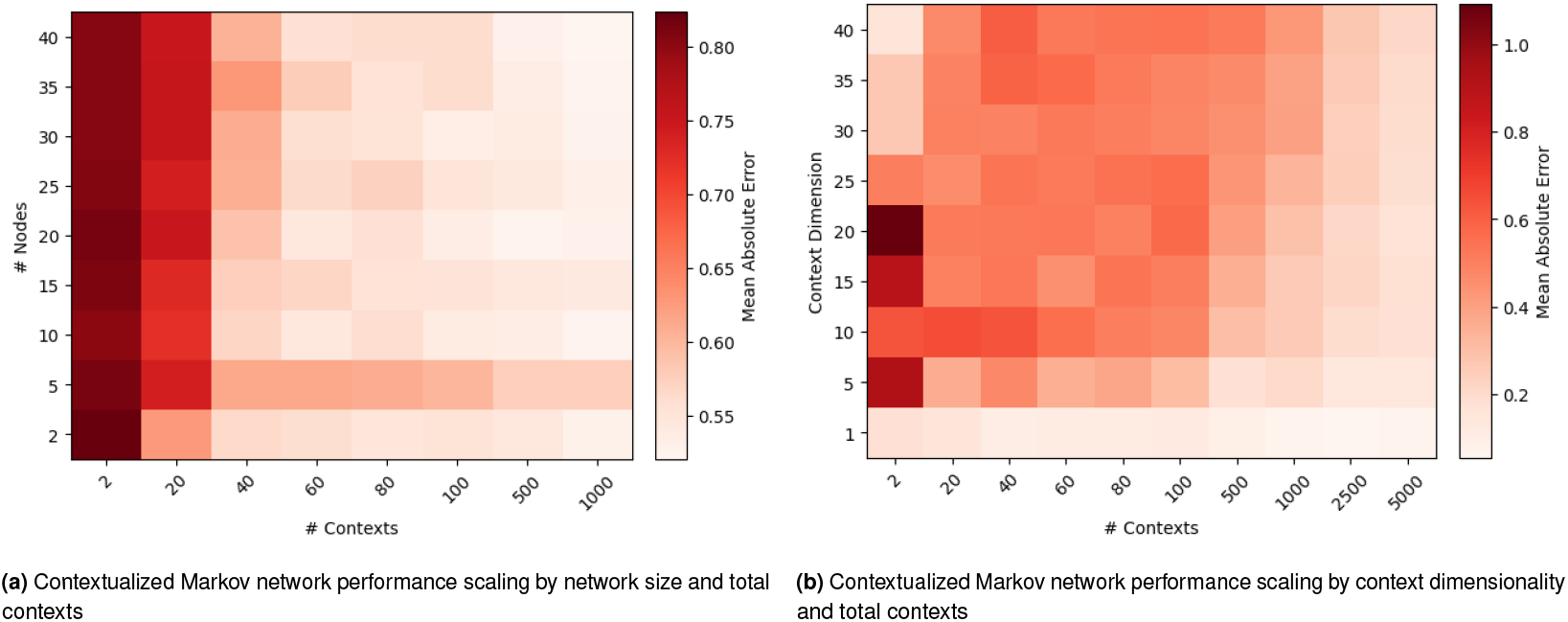
Performance of estimators on held-out data, measured across (a) network size and (b) context dimensionality (i.e. complexity of context dependence) measured as the Mean-absolute Error (MAE) between predicted and ground truth Markov networks, averaged over three bootstrapped runs.

**Fig. S3.**
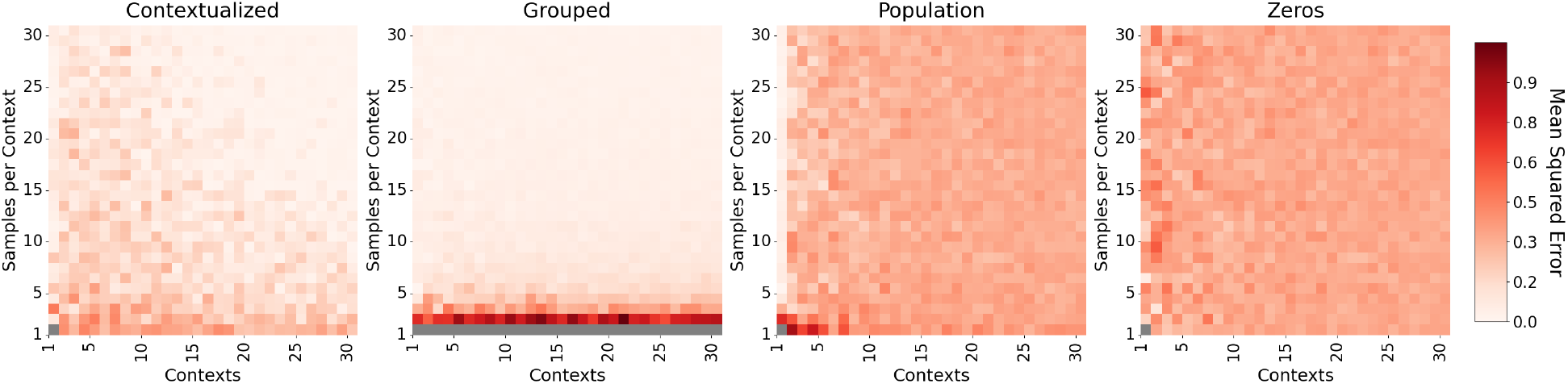
Performance of correlation network estimators in terms of predicted versus true network parameters, measured over samples per context task and total contexts. Mean-squared error (MSE) is between the predicted and ground truth correlation matrices defining the correlation networks, averaged over five bootstrapped runs. Population estimates a single model for all contexts, grouped estimates a model for each context separately. Contextualization is the only method to scale both horizontally and vertically.

**Fig. S4.**
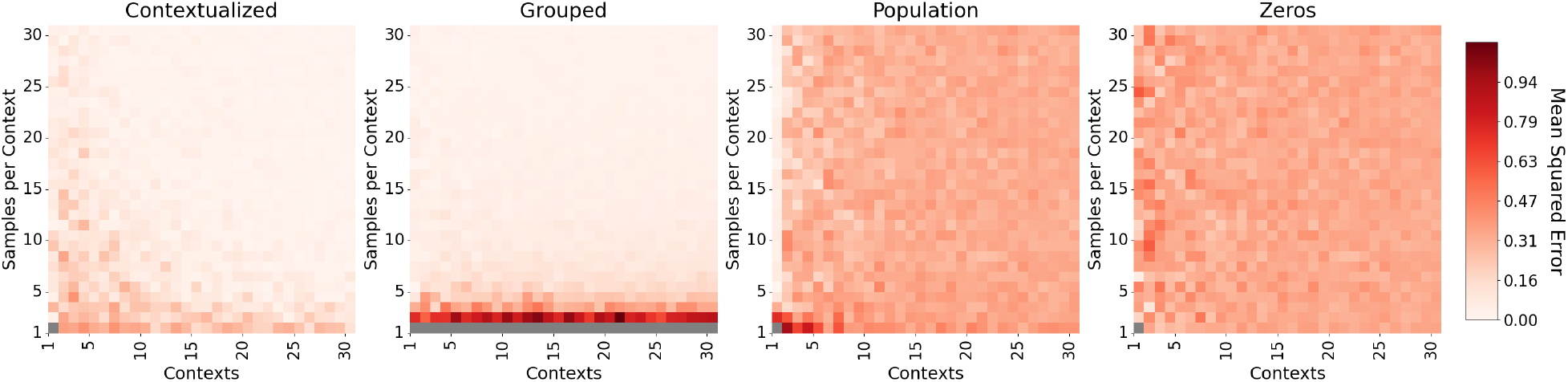
Performance of Markov network estimators in terms of predicted versus true network parameters, measured over samples per context task and total contexts. Mean-squared error (MSE) is between the predicted and ground truth precision matrices defining the Markov networks, averaged over five bootstrapped runs. Population estimates a single model for all contexts, grouped estimates a model for each context separately. Contextualization is the only method to scale both horizontally and vertically.

**Table S1.** Multivariate log-rank test comparison across different subtyping methods in terms of -log(p-value). Only samples shared between all datasets are used to control for power. – indicates no samples are shared, or subtypes do not exist for TCGA.

| Disease Type | Reference Subtypes | Expression Subtypes | Network Subtypes |
| --- | --- | --- | --- |
| Bladder Urothelial Carcinoma | 0.411 | 0.151 | 0.713 |
| Breast invasive carcinoma | 1.484 | 0.616 | 1.558 |
| Cholangiocarcinoma | – | – | – |
| Colon adenocarcinoma | 0.016 | 0.014 | 1.219 |
| Lymphoid Neoplasm Diffuse Large B-cell Lymphoma | – | – | – |
| Esophageal carcinoma | 0.044 | 0.884 | 0.049 |
| Glioblastoma multiforme | 1.644 | 0.958 | 0.101 |
| Head and neck squamous cell carcinoma | 1.209 | 0.312 | 3.465 |
| Kidney Chromophobe | 0.715 | 13.802 | 0.211 |
| Kidney renal clear cell carcinoma | 5.042 | 6.109 | 13.741 |
| Kidney renal papillary cell carcinoma | 14.538 | 10.582 | 15.205 |
| Brain Lower Grade Glioma | 48.338 | 33.438 | 49.681 |
| Liver hepatocellular carcinoma | 0.009 | 0.427 | 0.827 |
| Lung adenocarcinoma | 0.687 | 1.172 | 0.507 |
| Lung squamous cell carcinoma | 0.123 | 0.105 | 0.249 |
| Ovarian serous cystadenocarcinoma | 0.704 | 0.684 | 0.05 |
| Pancreatic adenocarcinoma | 0.439 | 1.104 | 1.494 |
| Prostate adenocarcinoma | – | – | – |
| Rectum adenocarcinoma | 0.221 | 0.203 | 0.117 |
| Skin cutaneous melanoma | – | – | – |
| Stomach adenocarcinoma | 0.044 | 1.117 | 0.575 |
| Thyroid adenocarcinoma | 0.298 | 0.164 | 2.104 |
| Uterine Corpus Endometrial Carcinoma | 6.937 | 3.343 | 7.07 |
| Uterine Carcinosarcoma | 0.319 | 0.023 | 0.048 |
| Uveal Melanoma | 4.838 | 2.589 | 3.565 |

**Table S2.** Minimum pairwise log-rank test comparison across different subtyping methods in terms of -log(p-value). Only samples shared between all datasets are used to control for power. – indicates no samples are shared, or subtypes do not exist for TCGA.

| Disease Type | Reference Subtypes | Expression Subtypes | Network Subtypes |
| --- | --- | --- | --- |
| Bladder Urothelial Carcinoma | 1.059 | 0.589 | 0.949 |
| Breast invasive carcinoma | 2.056 | 1.13 | 2.542 |
| Cholangiocarcinoma | – | – | – |
| Colon adenocarcinoma | 0.162 | 0.159 | 2.026 |
| Lymphoid Neoplasm Diffuse Large B-cell Lymphoma | – | – | – |
| Esophageal carcinoma | 0.323 | 1.598 | 0.284 |
| Glioblastoma multiforme | 1.631 | 1.509 | 0.63 |
| Head and Neck squamous cell carcinoma | 1.855 | 0.853 | 3.307 |
| Kidney Chromophobe | 0.715 | 13.802 | 0.211 |
| Kidney renal clear cell carcinoma | 5.61 | 5.247 | 14.82 |
| Kidney renal papillary cell carcinoma | 19.696 | 9.241 | 18.661 |
| Brain Lower Grade Glioma | 36.533 | 25.894 | 40.656 |
| Liver hepatocellular carcinoma | 0.105 | 0.619 | 1.263 |
| Lung adenocarcinoma | 1.67 | 2.29 | 1.198 |
| Lung squamous cell carcinoma | 0.614 | 0.417 | 0.625 |
| Ovarian serous cystadenocarcinoma | 1.414 | 1.325 | 0.287 |
| Pancreatic adenocarcinoma | 0.937 | 1.465 | 2.18 |
| Prostate adenocarcinoma | – | – | – |
| Rectum adenocarcinoma | 0.431 | 0.723 | 0.474 |
| Skin Cutaneous Melanoma | – | – | – |
| Stomach adenocarcinoma | 0.469 | 1.777 | 1.428 |
| Thyroid carcinoma | 0.837 | 0.831 | 3.242 |
| Uterine Corpus Endometrial Carcinoma | 5.555 | 3.319 | 7.42 |
| Uterine Carcinosarcoma | 0.319 | 0.023 | 0.048 |
| Uveal Melanoma | 5.076 | 2.61 | 4.536 |

**Table S3.** Concordance index of survival regressors for each tissue trained on different patient representations.

|  | Expression | Reference | Network |
| --- | --- | --- | --- |
| Bladder | 0.680 | 0.483 | 0.413 |
| Brain | 0.857 | 0.876 | 0.871 |
| Breast | 0.718 | 0.597 | 0.773 |
| Colorectal | 0.364 | 0.429 | 0.551 |
| Esophagus | 0.398 | 0.532 | 0.496 |
| Eye | 0.500 | 0.618 | 0.867 |
| Head and Neck | 0.558 | 0.587 | 0.572 |
| Kidney | 0.717 | 0.609 | 0.697 |
| Liver | 0.646 | 0.452 | 0.588 |
| Lung | 0.432 | 0.558 | 0.497 |
| Ovary | 0.412 | 0.591 | 0.536 |
| Pancreas | 0.468 | 0.634 | 0.542 |
| Thyroid | 0.475 | 0.413 | 0.580 |
| Skin | 0.424 | 0.436 | 0.333 |
| Stomach | 0.620 | 0.545 | 0.520 |
| Uterus | 0.767 | 0.742 | 0.632 |

**Fig. S5.**
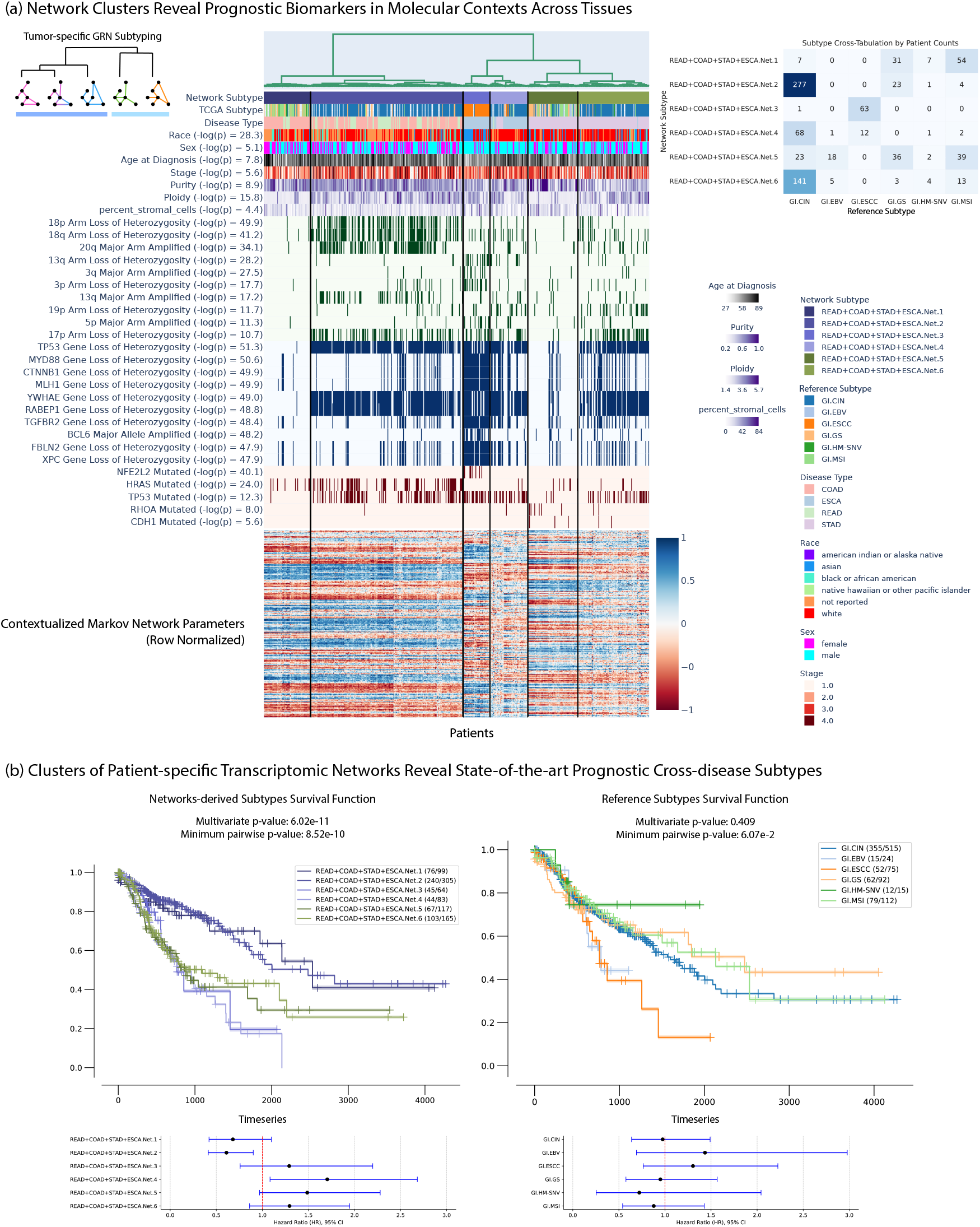
Exploration of cross-disease network subtypes for cancers of the GI tract, including rectum adenocarcinoma, colon adenocarcinoma, stomach adenocarcinoma, and esophageal carcinoma, looking at correlated clinical information, arm-level copy alterations, gene-level copy alterations, and gene-level single nucleotide variations. Reference subtypes from (17).

**Fig. S6.**
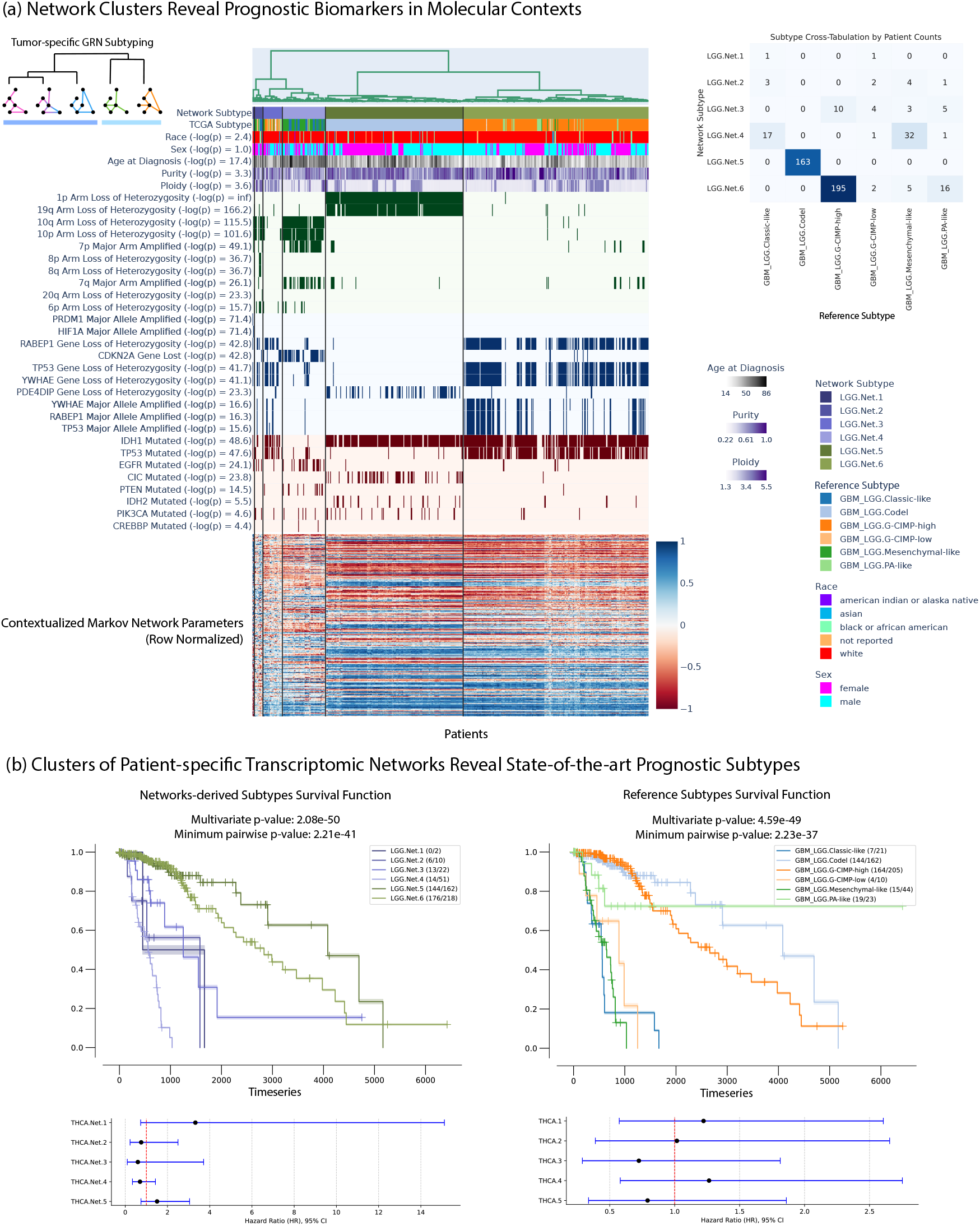
Exploration of Brain Lower Grade Glioma, looking at correlated clinical information, arm-level copy alterations, gene-level copy alterations, and gene-level single nucleotide variations. Reference subtypes from (18).

## SI Dataset S1 (data.tar.gz)

Pre-processed de-identified data from TCGA. https://zenodo.org/records/14885352/files/data.tar.gz

## SI Dataset S2 (results.tar.gz)

Pre-trained contextualized networks (correlation, Markov, neighborhood) for all 7997 patients with train/test split labels.

Includes subtyping plots for all 25 disease types with contextualized Markov networks. https://zenodo.org/records/14885352/files/results.tar.gz

## Footnotes

* https://www.cancer.gov/tcga

† www.cancer.gov/tcga

∗ www.cancer.gov/tcga

